# The Age of Selection-Duality Mutation under Fluctuating Selection among Individuals (FSI)

**DOI:** 10.64898/2026.01.30.701161

**Authors:** Xun Gu

## Abstract

Our recent work on molecular evolution and population genetics postulated that individuals with a specific mutation exhibit a fluctuation in fitness, short for FSI (fluctuating selection among individuals), whereas the fitness effect of wildtype remains a constant. An intriguing phenomenon called *selection-duality* emerges, that is, a slightly beneficial mutation could be a negative selection (the substitution rate less than the mutation rate). It appears that selection-duality is bounded by two bounds: the generic *neutrality* where the mutation is neutral by the means of fitness on average, and the *substitution neutrality* where the substitution rate equals to the mutation rate. In addition, the middle point of generic neutrality and substitution neutrality is called the FSI-*neutrality*. An important problem is about the age profile of allele frequency, i.e., the arising timing of a mutation whose frequency in the current population is given (*the allele-age problem* for short). Solving this problem under selection duality would help extend the standard coalescent theory that based on strict neutrality to a more general form under selection duality. In this paper, we studied the allele-age problem under selection-duality by the first arrival time approach and the mean age approach, respectively. Since the general solution of allele-age problem under selection duality is not available, we focused on solving the problem at the substitution neutrality (the up-bound of selection duality), the FSI-neutrality (the middle-point) and the generic neutrality (the low-bound), respectively. Our analysis results in an overall picture that the mean first-arrival age of a mutation at the substitution neutrality is theoretically identical to that at the FSI-neutrality, which is numerically close to that at the generic neutrality. For illustration, we calculated the mean age of nonsynonymous mutations in the human population and demonstrated that the estimated allele-age could be overestimated considerably when the effect of FSI was neglected.

## Introduction

In the population genetics theory of molecular evolution, the fixed view about the selection nature of a mutation is a fundamental assumption, which postulates that any single mutation has the same fitness effect among individuals with the same genotype (Crow and Kimura 1970; Kimura 1983). For instance, a neutral mutation is selectively neutral for all individuals who carry the mutation, and so forth a deleterious or beneficial mutation. By contrast, FSI, short for fluctuating selection among individuals, refers to the phenomenon when individuals with a specific mutation exhibit a broader phenotype variation, resulting in a fitness fluctuation, whereas the fitness of wildtype remains a constant.

The biological basis of the FSI of mutations can be well illustrated by the study of human geneticists have well-demonstrated that mutations frequently exhibit different effects on individuals (Riordan and Nadeau 2017; Eldar et al. 2009; Raj et al. 2010; Jensen et al. 2025). By the underlying mechanisms, FSI can be roughly classified into *genetic background* (Chandler et al. 2013; Mullis et al 2018), *stochastic gene expression* (Raj and van Oudenaarden 2007; Elowitz et al. 2002; Ozbudak et al. 2002; Maamar et al. 2007; Vu et al. 2015), *incomplete penetrance* (Khoury 1988; Eldar et al. 2009; Suel et al. 2007), as well as *the complexity of genotype-phenotype map* (Dowell et al. 2010; Lehner 2013; Taylor and Ehrenreich 2014). It appears that those categories are not mutually excluded (Raj et al. 2010).

The pattern of molecular evolution and population genetics of FSI has been studied recently (Gu 2025a; 2025b). Intriguingly, a novel phenomenon called ‘selection duality’ emerges from FSI: mutations that are statistically slightly beneficial are subject to a negative selection, which would merge to the conventional strict neutrality when FSI vanishes. Gu (2025b) showed that the substitution rate tends to inversely related to the log of effective population size (*N*_*e*_) when FSI is nontrivial, and developed a statistical procedure to predict the relative strength of FSI to the *N*_*e*_-genetic drift. Meanwhile, Gu (2025a) studied the population genetics of FSI, and in particular evaluated the effects of FSI on sequence divergence between species and genetic diversity within a population, revealing a provocative interpretation for the McDonald-Kreitman test that differs from the neutralist-view or the selectionist-view (Hahn 2008; Kern and Hahn 2018; Jensen et al. 2019; Munoz-Gomez et al. 2021; Gu 2021; Galtier 2024; de Jong et al. 2024).

An important problem is about the age profile of allele frequency, i.e., the arising timing of a mutation whose frequency in the current population is given. One may refer to *the allele-age problem* for short. Solving this problem under selection duality would help extend the standard coalescent theory that based on strict neutrality to a more general form under selection duality. In this paper, we studied the allele-age problem. Since the general solution under selection duality is not available, we focused on three special cases, that at the substitution neutrality (the up-bound of selection duality), the FSI-neutrality (the middle-point) and the generic neutrality (the low-bound), respectively. We implemented two approaches: the first arrival time refers to the expected generations required for the first arrival at the specified allele frequency, and the mean age approach to the average time it have reached. One may see Crow and Kimura (1970) or Ewens (2004) for the mathematical details. The aim of our research will focus on whether the analysis based on three type of neutrality can provide an overall picture about the allele-age in selection-duality. Our analysis is crucial for the study of coalescent theory. Although a deep coalescent analysis under FSI is out of the current scope, we speculate that the expectation of coalescent time under a constant population size would, when the sampling size is infinite, converge to the mean age of selection-duality mutation whose analytical form is given by the current study.

## Results

### The Wright-Fisher diffusion model under FSI

Consider a random mating population of a monoecioys diploid organism. In a finite population, each individual produces a large number of offspring and that exactly *N* of those survive to maturity. Let *a* and *A* be the mutant and wild-type alleles at a particular locus, respectively, whose fitness effects are additive. The FSI model postulates that the fitness effect of mutant *a* is fluctuating among individuals, whereas that of wild-type *A* remains a constant. Therefore, the relative fitness of genotype *AA, Aa*, or *aa* is, on average, given by 1, 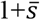 and 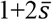, respectively; 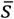 is called the mean of the selection coefficient (*s*) of mutant *a*. Let 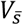 be the variance of 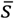 and *Var*(*s*) be the variance of *s*, respectively. Noting that the number of mutant *a* is 2*Nx*, where *x* is the frequency of mutant *a* in a generation, we obtain

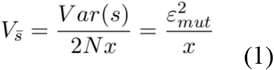

where 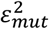 is called the FSI-coefficient; a large values means a strong FSI and *vice versa*; the subscript indicates the mutation-induced FSI.

Gu (2025a, 2025b) developed a diffusion model to study the Wright-Fisher model under FSI, under which the infinitesimal mean *μ*(*x*) and the infinitesimal variance *σ*^*2*^(*x*) are, respectively, given by

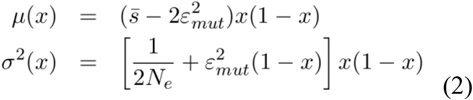

where *N*_*e*_ is the effective population size that inversely measures the strength of genetic drift with respect to the Wright’s sampling process in a finite population. Briefly speaking, *μ*(*x*) describes the determinative factors that may influence the gene frequency change, and *σ*^*2*^(*x*) describes the random effect of genetic drifts. In following-up analysis it is convenient to use the (adjusted) selection-FSI ratio (*ρ*) defined by

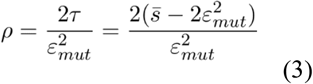

It appears that *ρ*>*0* indicates a positive selection, or *ρ*<*0* indicates a negative selection. Second, FSI emerges as a new resource of genetic drift, measured by the FSI-strength 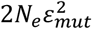 Reading the FSI-strength by 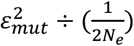, one may claim a dominant FSI-genetic drift when 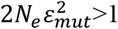, or a dominant *N*_*e*_-genetic drift when 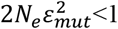. It is mathematically concise to use a relative measure of FSI-strength (*F*), as given by

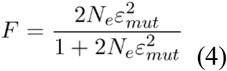

such that *F* increase from 0 at 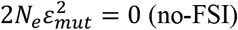, which approaches to 1 when 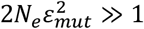.

### Substitution rate and emergence of selection duality

The substitution rate (*λ*) plays a central role in the theory of molecular evolution. Let *v* be the mutation rate and *N* be the census population size. From the view of population, the substitution rate can be defined by the amount of new mutations per generation (2*Nv*) multiplied by the fixation probability of a single mutation with the initial frequency of 1/(2*N*), based on the assumption of rare, single *de novo* mutation event. Let *u*(*p*) be the fixation probability of a mutation in a finite population, with the initial frequency *p*. Formally, the substitution rate can be written by 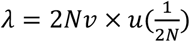 Gu (2025b) showed that

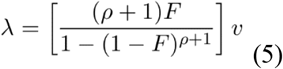

(One may also see Methods in details). It should be noticed that *λ*>*v* (the substitution rate greater than the mutation rate) when *ρ*>0, indicating a positive selection, whereas *λ*<*v* (the substitution rate less than the mutation rate) when *ρ*<0, indicating a negative selection. In the case of no-FSI, Eq.(5) is reduced to the well-known formula first reported by Kimura (1962).

Further analysis of Eq.(5) indicates that, when 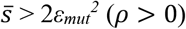 or 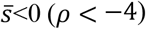, FSI only plays a marginal role in molecular evolution driven by a positive selection or a negative selection, respectively. However, between them, i.e., −4 < ρ < 0, an intriguing phenomenon called *selection-duality* emerges: a slightly beneficial mutation defined by

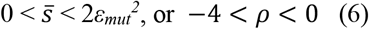

is subject to a negative selection (*λ*<*v* because of *ρ*<0). It appears that selection-duality defined by Eq.(6) is bounded by two types of neutralities. The low-bound is the generic neutrality 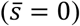, at which the mutation is neutral by the means of fitness. On the other hand, the up-bound of selection-duality is the substitution neutrality 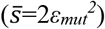, at which the substitution rate equals to the mutation rate (*λ*=*v*). The broadness of selection duality depends on the magnitude of *ε*_*mut*_^*2*^. Without FSI, i.e., *ε*_*mut*_^*2*^=0, the selection duality vanishes as the generic neutrality and the substitution neutrality merge onto the classical neutrality. In addition to those boundary neutralities, the middle-point of selection-duality at 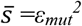 or *ρ*=-2, called FSI-neutrality, may play a pivotal role in the new theory of molecular evolution, as shown later.

### The mean age of a mutation with a given frequency: approximated by the first-arrival theory

Let 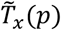 be the average number of generations until the mutant reaches frequency *x* for the first time starting a lower frequency *p*. As shown in Methods, *T*_*x*_(*p*) can be derived by the Wright-Fisher diffusion model (Kimura and Ohta 1973). It should be noticed that, as 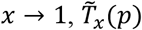 equals to the mean fixation time of a mutation *T*_*fix*_(*p*), given the initial frequency *p* (Kimura and Ohta 1969). We are particularly interested in the average number of generations until the mutant reaches frequency *x* for the first time since its single origin with the allele frequency *p*=1/(2*N*), where *N* is the census population size. Therefore, the age distribution of a mutant since its origin can be approximated by *p* → 0, that is,

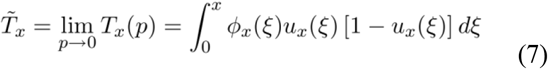

There are two functions in Eq.(7). The first one is *u*_*x*_(*ξ*), the probability of a mutant to first reach frequency *x* before being lost, i.e., reaching the zero-boundary, given the initial frequency *ξ*, where the condition 0 < ξ ≤ *x* holds. It can be calculated by

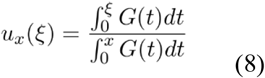

where G(ξ) in Eq.(8) is defined by

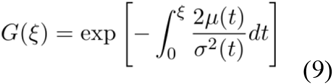

Apparently, as *x*=1, *u*_*x*_(*ξ*) becomes the well-known fixation probability. The second function is *ϕ*_*x*_(*ξ*), the sojourn time in the interval of (0, *x*], given the initial frequency *ξ* that satisfies 0 < ξ ≤ *x*, as given by

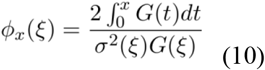

The solution of 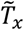 by Eq.(7) is mathematically complex; there is no analytical solution in general. Nevertheless, we are interested the (first-arrival) age of mutations at some special case of selection duality, as shown below.

### 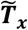 at substitution neutrality identical to that at FSI-neutrality but differs marginally from that at generic neutrality

As discussed above, the selection-duality defined by Eq.(6) is bounded by two types of neutralities. The low-bound is the generic neutrality (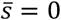, or *ρ* = −4), at which the mutation is neutral by the means of fitness on average. Meanwhile, the up-bound of selection-duality is the substitution neutrality (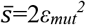, or *ρ* = 0), at which the substitution rate of a mutation equals to the mutation rate (*λ*=*v*). In addition to those boundary neutralities, the middle-point of selection-duality is given by 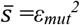 or *ρ*=-2, called FSI-neutrality. Note that the analytical solution of 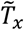 is not available for the range of selection-duality. Instead, we try to analyze 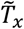 at each of three neutralities respectively so that by putting together, one can provide an overall pattern of 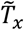 under selection duality.

Intriguingly, we found a surprising result that 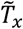 at the substitution neutrality is identical to that at the FSI-neutrality. While the detailed derivation can be found in Methods, a brief argument is presented here. We first calculate the production of *ϕ*_*x*_(*ξ*)*u*_*x*_(*ξ*)[1 − *u*_*x*_(*ξ*)]. It has been shown that at either substitution neutrality (*ρ* = 0) or at FSI-neutrality (*ρ* = −2), this product is given by

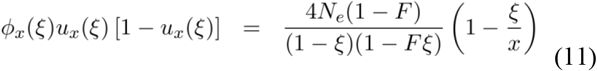

According to Eq.(7), one can immediately conclude that the (first-arrival) age of a mutation, 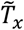 at the substitution neutrality and that at the FSI-neutrality are the same, which is given by

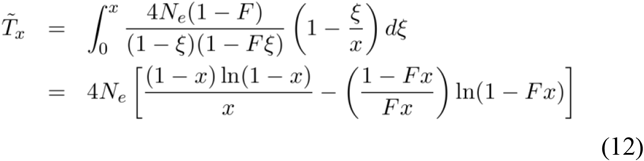

It appears that 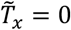 when *x* = 0, which means that a mutation with very low frequency indicates a very recent origin. On the other hand, we have 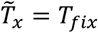 when *x* = 1, i.e., the mean fixation time, as given by

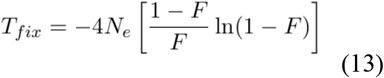

which is further reduced to 4*N*_*e*_ when *F* = 0, i.e., strictly neutral mutations without FSI. Fig.1 shows the plotting of the (first-arrival) age distribution 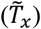 against the allele frequency (*x*) at FSI-neutrality. While 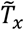 increases symbolically with the increase of *x*, the effect of FSI is overall to reduce the age of mutations.

**Fig.1.**
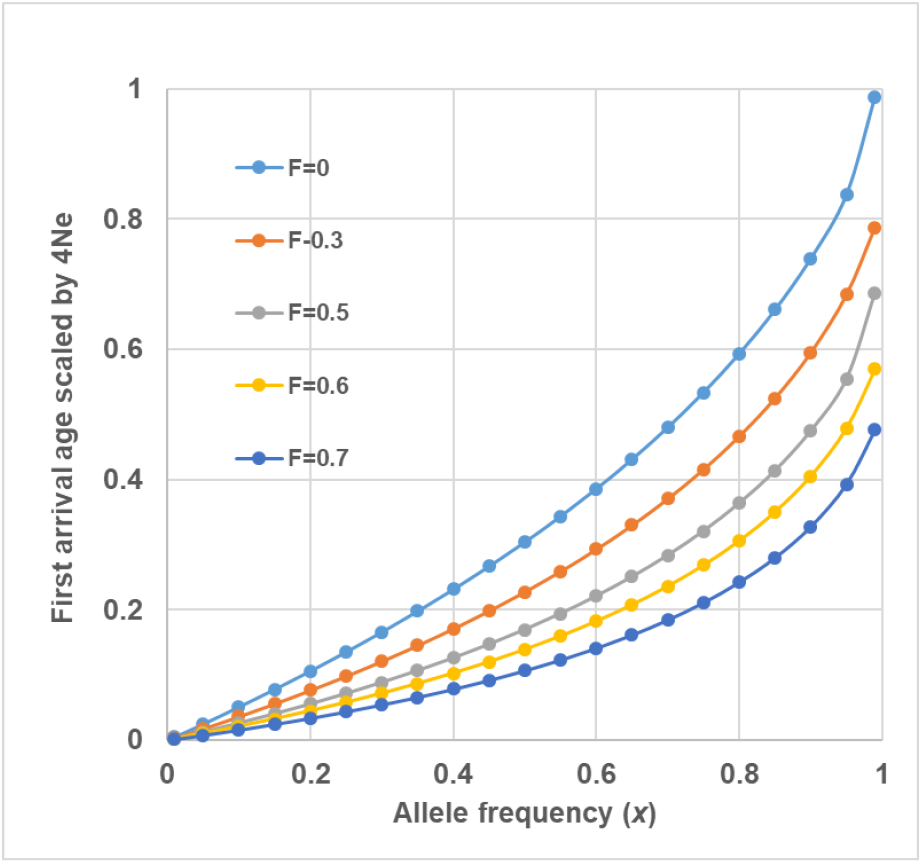
The first-arrival age of alleles plotting against the allele frequency under different levels of FSI

One may wonder whether 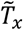 at the generic neutrality is also the same. To this end, we examined the product of *ϕ*_*x*_(*ξ*)*u*_*x*_(*ξ*)[1 − *u*_*x*_(*ξ*)] by Eq.(M6) with *ρ*=-2. As shown by Eq.(M10), one may concluded that the first-arrival age of a mutation at the generic neutrality differs from that at the substitution neutrality or the FSI-neutrality.

Moreover, 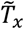 at the generic neutrality given by Eq.(M11) is analytical but tedious. Nevertheless, a direct numerical integral analysis showed that the difference of 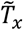 between the generic neutrality and the substitution/FSI-neutrality should be marginal (not shown).

### The mean age of a substitution-neutrality mutation in a finite population

Another method to evaluate the age distribution of mutations is the mean generations since an allele (that now has intermediate frequency *x*) had a lower frequency *p*, i.e., *p* ≤ *x*. With some technical modifications and refinements, I follow the approach proposed by Kimura and Ohta (1973) to derive the mean age of a mutation in a finite population in the case of substitution neutrality.

Let *ϕ*(*x*, *p*; *t*) be the probabilistic density of allele frequency (*x*) at time *t*, given by the initial frequency *p*. It follows that, *T*_*i*_(*x*), the *i*-th moment of *t*, (*i*=0, 1, 2, …) of a mutation with the gene frequency (*x*), given the initial frequency *p*, is determined by

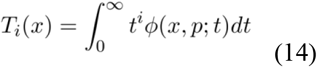

Therefore, the mean age of mutation with allele frequency (*x*) can be calculated by

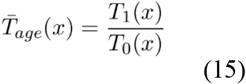

Let 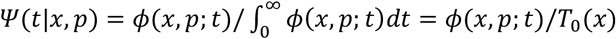 be the time (or allele age) -distribution conditional of the allele frequency (*x*) and the initial frequency (*p*). It follows that Eq.(15) can be written as 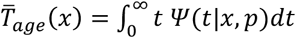, which can be intuitively interpreted in a conversional way.

Kimura and Ohta (1973) showed that *T*_*0*_(*x*) and *T*_*1*_(*x*) satisfy the following ordinary differential equations

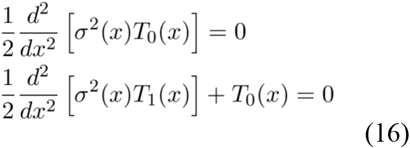

Note that Eq.(16) must satisfy the following constraints: *T*_*0*_(*x*)< ∞ for the regularity of probabilistic density, and *T*_*1*_(*x*) < ∞ for *p* ≤ *x* ≤ 1, which means that a mutant with the initial frequency *p* can reach the frequency of *x* in a finite time. As shown by Methods in details, we obtain

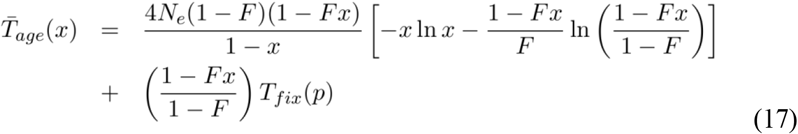

In particular, we are interested in 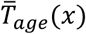 in the case of very low initial frequency such that *p* → 0; in this case *T*_*fix*_ = *T*_*fix*_(0) is given by Eq.(13). One can further verify that in the case of no-FSI, i.e., *F*=0, Eq.(17) is reduced to the well-known formula

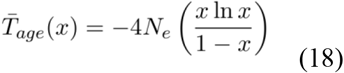

### Case study: the mean age of nonsynonymous mutations in the human population

We utilized the human population genetics data of Fu et al. (2013) to carry out a preliminary analysis. Fu et al. (2013) re-sequenced about fifteen thousand genes over six thousand individuals of European American and African American ancestry and inferred the age of over one million autosomal single nucleotide variants (SNVs). Among different types of variants they studies, we focused on nonsynonymous mutation in protein-coding genes, because our previous study (Gu 2025a) has shown an intermediate FSI (*F* ≈ 0.5) in those mutations in the human population. It was estimated (Fu et al. 2013) that the average age of nonsynonymous mutations was about 2.1 × 10^4^ years (European American) or about 3.1 × 10^4^ years (African American). Those estimates were obtained under the assumption of *F* = 0. By Eq.(17) and Eq.(18), we re-estimated those ages under *F* = 0.5 and obtained the average ages was about 1.0 × 10^4^ years (European American) or about 1.4 × 10^4^ years (African American). It appears that the average age of nonsynonymous mutations could be considerably overestimated if FSI is not considered.

## Discussion

### The allele age of selection-duality mutation

In this work we addressed the allele-age problem under the Wright-Fisher model of FSI (fluctuating selection among individuals). Two measures are studied. The first one is the mean first-arrival age 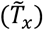 of a mutation to the frequency *x*. We are particularly interested in the case of selection duality, where a mutation that is slightly beneficial on average is actually subject to a negative selection. Since the general solution of 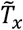 in the range of selection duality is not available, we focused on 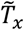 at the substitution neutrality (the up-bound of selection duality), the FSI-neutrality (the middle-point) and the generic neutrality (the low-bound), respectively. The overall picture we attempted to provide is as follows

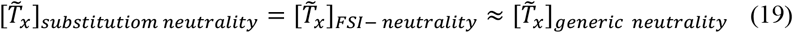

That is, the mean first-arrival age 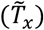 of a mutation at the substitution neutrality is theoretically identical to that at the FSI-neutrality, which is numerically close to that at the generic neutrality. Tentatively, we propose that the age profile of allele frequency may be virtually universal in selection-duality.

On the other hand, we used the method of Kimura and Ohta (1973) to derive the mean age of a mutation with frequency *x* at the substitution neutrality, 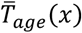 . One may envisage that 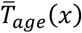 is also universal in selection-duality. While this claim is rational, the method we used here cannot be applied to the case of FSI-neutrality nor the generic-neutrality. It remains further study to test whether a relationship similar to Eq.(19) also holds for 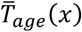 . One possible approach to solving this issue is to invoke the method developed by Maruyama (1974) which is mathematically sophisticated.

### The coalescent theory of allele age

Coalescent theory is a powerful framework in population genetics that simulates the ancestry of genes backward in time, tracing sampled DNA lineages back until they merge (coalesce) into a single common ancestor. There is an intrinsic relationship between coalescence and the mean age of a mutant given the allele frequency. Let *T*_*n*,*b*_ denote the age of a mutant having *b* copies in a sample of *n* genes (0 < *b* < *n*). Griffiths and Tavaré (1998) have derived the general formulas for the mean and variance of *T*_*n*,*b*_, respectively. In the case of constant population, for instance, the expectation of *T*_*n*,*b*_ is given by

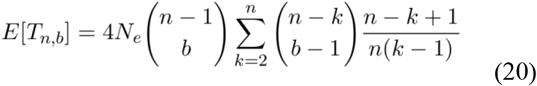

Griffiths and Tavaré (1998) showed that, when the sample size approaches to infinite, i.e., *n* → ∞, the sample frequency *b*/*n* → *x*, and so the expected *T*_*n*,*b*_ approaches to the mean age of a strictly neutral mutant with frequency (*x*), that is,

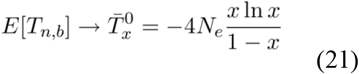

which was first derived by Kimura and Ohta (1973), also shown by Eq.(18).

An intriguing problem is how to extend the coalescent theory to the case of FSI. Our goal is to formulate the coalescent framework of selection-duality mutant under FSI such that the expected *T*_*n*,*b*_, denoted by 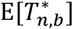, converges to 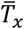 as the sample size approaches to infinite. One may speculate that it is technically very difficult and some approximations must be made. Moreover, a concept clarification is more challenging:there are three types of neutrality under FSI: substitution neutrality, FSI-neutrality and generic neutrality, which reveal distinct population genetics features such as sampling property (Sawyer and Hartl, 1992) ; one see Gu (2025b) for a detailed analysis. At the current stage, it remains unclear for their relationship with the coalescent neutrality. As the coalescent analysis treats genealogies as random processes rather than fixed trees, it seems difficult, under FSI, to derive an elegant probabilistic framework to understand how present-day genetic diversity arose from past events such as demographic histories. The impact of FSI on coalescent-based inference will definitely be the goal of future study.

## Methods

### Age of a mutation with a given allele frequency: approximated by the first-arrival theory

Let *µ*(*x*) and *σ*^2^(*x*) be, respectively, the mean and the variance of the change in one generation of the frequency of a mutant allele having frequency *x*. This diffusion model assumes that the stochastic process of change in gene frequency is time homogenous, that is, the mean selection coefficient of the mutant remains constant with time while it may fluctuate among individuals. The average number of generations until the mutant reaches frequency *x* for the first time starting a lower frequency *p*, denoted by 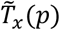, can be derived by the diffusion method. Let *u*_*x*_(*ξ*) be the probability of a mutant to first reach frequency *x* before being lost, i.e., reaching the zero-boundary, given the initial frequency *ξ*, where the condition 0 < ξ ≤ *x* holds. It can be calculated by

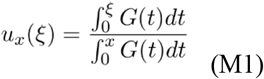

where G(ξ) is defined by

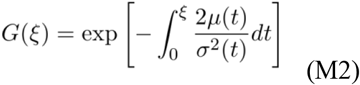

Meanwhile, let *ϕ*_*x*_(*ξ*) be the sojourn time in the interval of (0, *x*], given the initial frequency *ξ* that satisfies 0 < ξ ≤ *x*, as given by

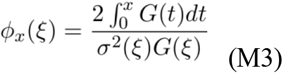

It follows that the average number of generations until the mutant reaches frequency *x* for the first time starting a lower frequency *p* is given by

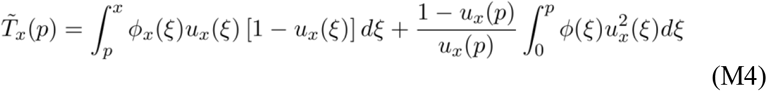

The first term on the right hand of Eq.(M4) represents the average sojourn time of the mutant with the frequency between *p* and *x*, while the second term presents that of the mutation with the frequency between 0 and *p*. It should be noticed that *T*_*fix*_(*p*), the mean fixation time of a mutation, given the initial frequency *p* (Kimura and Ohta 1969), is a special case of *T*_*x*_(*p*) as *x* → 1, that is,

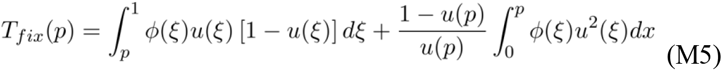

where *u*_*x*_(ξ) → *u*(ξ) and *ϕ*_*x*_(*ξ*) → *ϕ*(*ξ*). It appears that the average number of generations until the mutant reaches frequency *x* for the first time since its single origin can be calculated by allowing the allele frequency *p*=1/(2*N*), where *N* is the census population size. In this case, the age distribution of a mutant since its origin can be approximated by *p* → 0. One can show that the second term on the right hand of Eq.(M5) approaches to 0, leading to Eq.(7).

### (First-arrival) age of selection duality mutations

#### The general formula

According to Eqs.(2-4), one can derive the following function under the FSI model, that is,

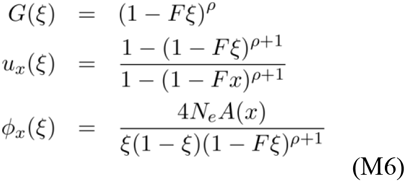

where function *A*(*x*) is given by

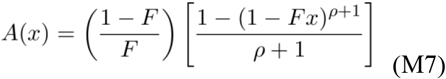

By some calculus analyses, one can show that the analytical solution of Eq.(7) is available only when ρ is an integer (positively or negatively or zero) except for ρ=-1. Since we are interested in the scenario of selection duality defined by the range of −4 ≤ ρ ≤ 0, we try to derive 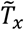 at the FSI neutrality (ρ = −2), the substitution neutrality (ρ = 0), or generic neutrality (ρ = −4), which, together, provide an overall pattern of the age distribution of selection duality mutations.

#### Age of mutation at FSI neutrality 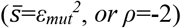

By Eq.(M6), three functions, *u*_*x*_(*ξ*), *ϕ*_*x*_(*ξ*) and *A*(*x*) under *ρ*=-2 are, respectively, are given by

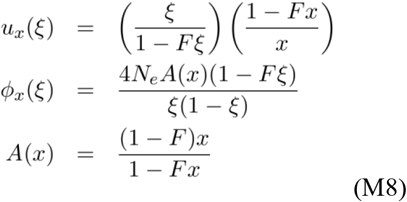

and so the product of those variables is then given by Eq.(11). We thus derive the (first-arrival) age distribution of mutations at FSI-neutrality by Eq.(12).

#### Age of substitution neutrality mutation 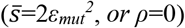

In this case of substitution neutrality (*ρ* = 0), we have the expressions of three variables, *u*_*x*_(*ξ*), *ϕ*_*x*_(*ξ*) and *A*(*x*) as follows

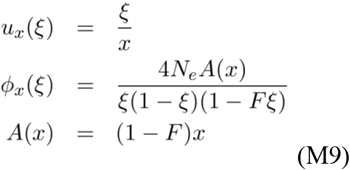

In spite that each function in Eq.(M9) differs from the corresponding ones in Eq.(M8), we find that the product, *ϕ*_*x*_(*ξ*)*u*_*x*_(*ξ*)[1 − *u*_*x*_(*ξ*)] based on Eq.(M9) is precisely identical to Eq.(M8), the same product in the case of FSI-neutrality. In other words, the (first-arrival) age distribution of mutation at the substitution neutrality is identical to that at the FSI-neutrality.

#### Age of generic mutation 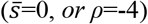

Plugging *ρ* = −4 into Eq.(M6), we have

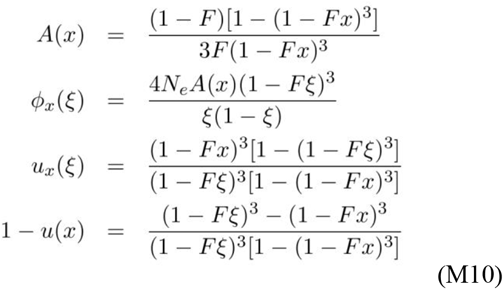

Eq.(M10) shows that the first-arrival age of a mutation at the generic neutrality differs from that at the substitution neutrality or the FSI-neutrality. The age distribution of mutations at generic neutrality is then given by

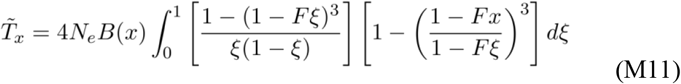

where *B*(*x*) is for

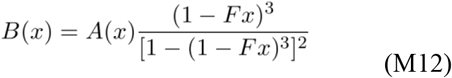

The result of Eq.(M11) is tedious in algebra; yet a numerical integral analysis based on Eq.(M11) is straightforward. Besides, for the purpose of comparison, one may calculate the age distribution of mutations at a negative selection-duality 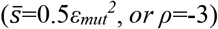 analytically, with the following specifications

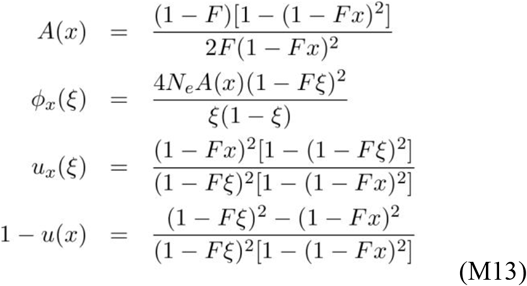

### The mean age of a substitution-neutrality mutation

We first formulate this problem briefly. Let *T*_*i*_(*x*) be the *i*-th moment of *t*, (*i*=0, 1, 2, …) of a mutation with the gene frequency (*x*), given the initial frequency *p*, as given by Eq.(14). It follows that the mean age of mutation with allele frequency (*x*) can be calculated by

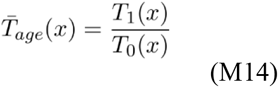

Kimura and Ohta (1973) showed that *T*_*0*_(*x*) and *T*_*1*_(*x*) satisfy the following ordinary differential equations

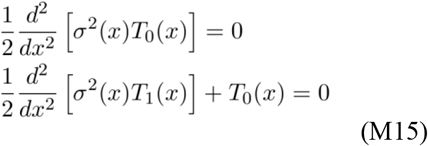

which satisfies the following constraints: *T*_*0*_(*x*)< ∞, and *T*_*1*_(*x*) < ∞ for *p* ≤ *x* ≤ 1.

We first consider *T*_0_ (*x*). By integrating twice the first differential equation of Eq.(M15), and rewriting *σ*^2^(*x*) by

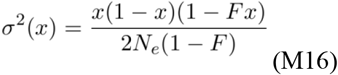

we obtain

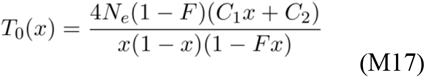

where *C*_*1*_ and *C*_*2*_ are two arbitrary constants. The constraint of *T*_*0*_(*x*)< ∞ for *p* ≤ *x* ≤ 1 implies that the term *C*_1_*x* + *C*_2_ in the numerator should be canceled with the term 1 − *x* in the denominator. Without loss of generality, one may choose *C*_2_ = −*C*_1_ = *A*/[4*N*_*e*_(1 − *F*)], where *A* is an arbitrary constant, resulting in

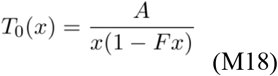

Therefore, the differential equation of *T*_*1*_(*x*) in Eq.(M15) can be specified as follows

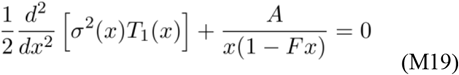

Integrating this equation twice with respect to *x*, we obtain

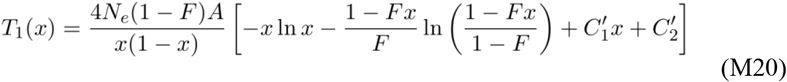

Where 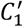 and 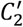 are arbitrary constants that can be determined as follows. The constraint of *T*_*1*_(*x*)< ∞ for *p* ≤ *x* ≤ 1 implies that the term *C*′_1_*x* + *C*′_2_ should be canceled with the term 1 − *x* in the denominator. Without loss of generality, one may choose *C*′_2_ = −*C*′_1_ = *B*, resulting in

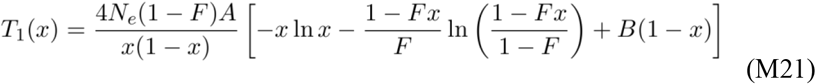

It follows that, by Eq.(M18), the mean age of mutation under the substitution neutrality can be expressed as follows

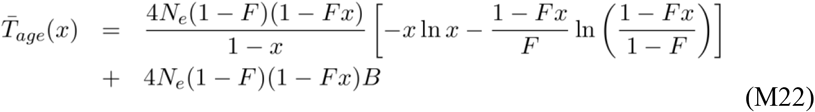

which is independent of constant *A*. The arbitrary constant *B* can be determined by the boundary when the allele frequency *x* approaches unity: *T*_*age*_ (*x*) should approach the average number of generations until fixation (Kimura and Ohta 1969), that is,

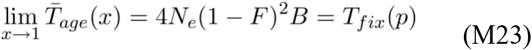

Replacing *B* in Eq.(M22) by Eq.(M23), we obtain

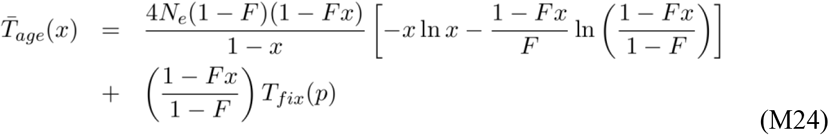

## Notes

### Competing Interest Statement

The authors have declared no competing interest.

